# Comparative analysis of Endoxifen, Tamoxifen and Fulvestrant: A Bioinformatics Approach to Uncover Mechanisms of Action in Breast Cancer

**DOI:** 10.1101/2024.10.02.616224

**Authors:** H. Lawrence Remmel, Sandra S. Hammer, Harjinder Singh, Anastasia Shneyderman, Alexander Veviorskiy, Khadija M. Alawi, Mikhail Korzinkin, Alex Zhavoronkov, Steven C. Quay

**Affiliations:** Atossa Therapeutics Inc., 10202 5th Ave NE Suite 200 Seattle, WA 98125; Department of Pathology, University Medical Center Utrecht, Utrecht, The Netherlands; Children’s Cancer and Blood Foundation Laboratories, Departments of Pediatrics, and Cell and Developmental Biology, Drukier Institute for Children’s Health, Meyer Cancer Center, Weill Cornell Medicine, New York, NY, USA; Insilico Medicine Hong Kong Ltd., Hong Kong Science and Technology Park, Hong Kong, Hong Kong SAR, China; Insilico Medicine AI Limited, Abu Dhabi, UAE; Insilico Medicine Canada Inc., 3710-1250 René-Lévesque Blvd. W, Montreal, Quebec, Canada H3B 4W8; Buck Institute for Research on Aging, Novato, California 94945, United States

**Author notes:** **Corresponding author:** H. Lawrence Remmel.

## Abstract

Breast cancer remains a significant health challenge, with estrogen receptor positive (ER+) subtypes being particularly prevalent forms of breast cancer. Current anti-estrogen therapies, such as tamoxifen and fulvestrant, have limitations, including partial agonist activity and resistance development, which evidence the need for more potent alternatives. Endoxifen, a metabolite of tamoxifen, has emerged as a promising breast cancer therapeutic candidate due to its superior anti-estrogenic effects and side effect profile. The omics signatures for endoxifen, tamoxifen and fulvestrant, obtained from publicly available datasets, were aggregated and harmonized by means of the PandaOmics platform, a commercially available target-discovery platform using multiple AI engines including generative pretrained transformers. Pathway enrichment analyses provided insight into these agents’ mechanisms of action (MOA) in breast cancer. The analyses revealed unexpected variances in several key pathways from expected interactions via estrogen-dependent and independent effects. All three drugs downregulated estrogen signaling and cell cycle-related pathways, such as E2F targets, G2-M checkpoints, Myc targets, and mitotic spindle, and stimulated apoptosis. Fulvestrant and tamoxifen activated pro-inflammatory and immune pathways and perturbed epithelial-mesenchymal transition (EMT). Endoxifen perturbed the PI3K/Akt/mTORC1 pathway, pursuant to distinct molecular mechanisms compared to its parent compound, tamoxifen, and fulvestrant. In summary, advanced AI-driven methodologies demonstrate the capacity to analyze multi-omics data in a comparative way to advance the understanding of endocrine therapy mechanisms in breast cancer. This insight into the distinct effects of endoxifen, tamoxifen, and fulvestrant may aid in selecting the most effective therapies for specific indications and in identifying drug-specific biomarkers.

## Introduction

Breast cancer remains a significant global health challenge, with high incidence and mortality rates impacting healthcare systems worldwide. The majority of breast cancers are estrogen receptor-positive (ER+), relying on estrogen signaling for growth and proliferation. Targeting estrogen signaling pathways with endocrine therapies has been a cornerstone of breast cancer therapy (Hanker, Sudhan, and Arteaga 2020). However, current treatments for hormone receptor-positive breast cancer with endocrine therapy face limitations due to side effects, resistance development, and varying efficacy across different subtypes of breast cancer. Tamoxifen, a selective estrogen receptor modulator (SERM), exhibits both estrogen agonist and antagonist properties, which can lead to adverse events such as tumor flare and endometrial cancer in addition to its antitumor effects (Veldhuis and Santen 1979; Emons, Mustea, and Tempfer 2020). Fulvestrant, a pure anti-estrogen and a selective estrogen receptor degrader (SERD), degrades the ER and inhibits its signaling, but its clinical efficacy is sometimes limited by the development of resistance (Giessrigl et al. 2013). Endoxifen (a SERM) is one of the most potent and bioactive metabolites of tamoxifen, produced through the metabolism of tamoxifen by the enzyme CYP2D6. Emerging studies have focused on endoxifen due to its potent anti-estrogenic effects, which are superior to its parent compound, tamoxifen, highlighting its potential in offering a more effective treatment option for patients who are poor metabolizers of tamoxifen. Clinical trials of endoxifen have demonstrated its efficacy and safety, suggesting that it could be a valuable alternative in the treatment of ER+ breast cancer (Jayaraman et al. 2021). Understanding the various effects of available hormonal therapies should optimize treatment selection for breast cancer patients.

To determine the various effects of available hormonal therapies, multi-omic publicly available datasets were aggregated and harmonized by means of the PandaOmics platform to generate a signature for each of endoxifen, tamoxifen and fulvestrant (Ren et al. 2024; Kamya et al. 2024). The integration of bioinformatics and multi-omics analyses generated data that clarified the molecular mechanisms of endoxifen, tamoxifen and fulvestrant and the shared or varying effects of these three anti-estrogens. The data obtained confirmed that all three drugs effectively inhibit estrogen signaling and disrupt the cell cycle in breast carcinoma, and confirmed as well that the other effects of each agent differ. Tamoxifen and fulvestrant specifically induce pro-inflammatory responses and disrupt epithelial-mesenchymal transition (EMT), whereas endoxifen is associated with the regulation of the PI3K/Akt/mTORC1 pathway. These data provide new insights in understanding of the two SERMs’ and the SERD’s mechanisms applicable to breast cancer.

## Materials and Methods

### Data Collection and Integration Using PandaOmics AI Platform

Gene expression data obtained from Gene Expression Omnibus (Barrett et al. 2013), ArrayExpress (Parkinson et al. 2007), and PRIDE (Perez-Riverol et al. 2022) were collected in PandaOmics, an AI-driven target discovery platform (Kamya et al. 2024), to create meta-analyses. For endoxifen (four datasets), tamoxifen (thirteen datasets), fulvestrant (twenty-one datasets) and estrogen (twelve datasets), the primary data were derived from MCF-7 cells, a widely used model in breast cancer research (Booms, Coetzee, and Pierce 2019), treated under different conditions (see supplementary table 1).

#### Breast carcinoma

a pre-prepared, manually curated meta-analysis of breast carcinoma (Experimental Factor Ontology: EFO_0000305 (Malone et al. 2010)) is available in PandaOmics. This meta-analysis is composed of 34 multi-omic datasets (RNA-seq, microarray, proteomics and methylation) with 4019 total samples. The dataset information is provided in supplementary table 1. All omics datasets were pre-processed according to the PandaOmics pipeline, which automatically defines data type and normalizes the data for further analysis (Kamya et al. 2024).

### Data analysis

#### Differential expression analysis and combined log-fold changes

Differential expression analysis was performed in PandaOmics using the *limma* R package. Each dataset has been processed according to *limma* standard protocols. Obtained gene-wise p-values were corrected by the Benjamini-Hochberg procedure. Combined log-fold changes (LFC) between transcriptomics and proteomics datasets were calculated by PandaOmics using the ‘Expression’ feature. This feature calculates combined LFC values and Q-values across all datasets used for the analysis, using minmax normalization for LFC values and Stouffer’s method combining p-values with further false discovery rate (FDR) correction. All differentially expressed genes (DEGs) from the respective meta-analyses were downloaded and used to generate the final drug specific gene expression signatures.

#### Pathway enrichment analysis

Pathway enrichment analysis was performed with the GSEApy package (Fang, Liu, and Peltz 2023) using the enrichr() function according to standard protocols. Differentially expressed genes were used as a gene list for pathway enrichment analysis. MsigDB database was selected as the gene set from GSEApy internal library for signaling pathway enrichment analysis.

#### Estrogen-dependent and Independent Effects

An estrogen-dependent gene signature was generated using available omics data. Genes that were differentially expressed following estrogen treatment were classified as estrogen-dependent, while all other genes were deemed estrogen-independent. Subsequently, gene set enrichment analysis (GSEA) analyses were conducted on both the estrogen-dependent and estrogen-independent genes, as previously described.

#### MOA Analysis in Breast Cancer

The molecular signature for breast cancer was developed using relevant data on the PandaOmics platform. To identify the molecular effects of the three anti-estrogens in breast cancer, individual drug signatures were intersected with the disease signature. This process identified genes that are upregulated in breast cancer but downregulated after treatment, and conversely, genes that are downregulated in cancer compared to healthy tissue but upregulated following treatment. GSEA analyses were performed on the genes in the intersection as previously described.

#### Paper draft preparation

The draft for the current paper was generated using the Draft Outline Research Assistant (DORA, https://dora.insilico.com/), Insilico Medicine’s Large Language Model (LLM) based text drafting assistant. DORA is designed to streamline the process of publication drafting, making it faster and simpler. The text generation is curated by over 30 AI agents, powered by LLMs, and integrated with internal and other curated databases, to assist in generating high-quality scientific document drafts. Each agent employs Retrieval-Augmented Generation (RAG) to perform comprehensive data collection and analysis, reduce the probability of poorly founded analyses, and provide relevant PubMed links to make the generation of the article more transparent. The draft was manually curated, commented on, extended and reviewed, and appropriate citations added, by all authors.

## Results

### Analysis of individual gene expression signatures of tamoxifen, endoxifen and fulvestrant

In order to study molecular changes after tamoxifen, endoxifen and fulvestrant treatment, the most complete publicly available omics data for the three anti-estrogen drugs were collected. For each drug the gene expression profile of MCF7 cells exposed to a drug was compared relative to control, and individual comparisons were combined into meta-analyses. Combined results were used to generate the final drug specific gene expression signatures. GSEA analyses were performed to identify signaling pathways that are perturbed by each drug (Figure 1A).

**Figure 1.**
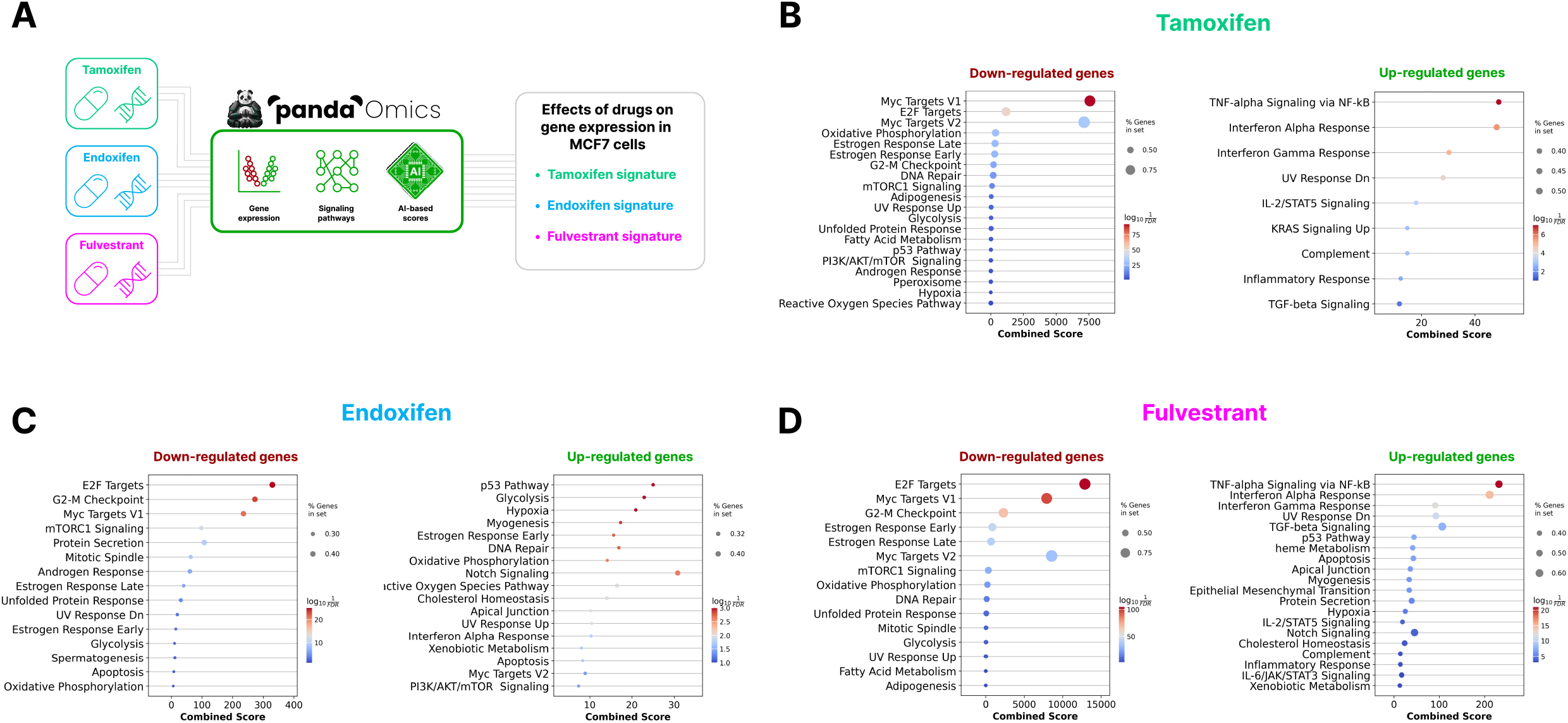
A) Drug analysis pipeline. The most complete publicly available omics data for the three antiestrogen drugs, tamoxifen, endoxifen and fulvestrant, were collected. Data were integrated in PandaOmics, and the final drug specific gene expression signatures were generated. Figure 1 reflects the results of gene set enrichment analysis (GSEA) conducted on genes that are downregulated or upregulated after tamoxifen (B), endoxifen (C), or fulvestrant (D) treatment in MCF7 cells.

### Tamoxifen gene expression signature

(Figure 1B) was characterized by downregulated genes that enrich early and late hallmarks of estrogen response. This characterization may be considered as a confirmation of anti-estrogenic effects of a SERM. Also, cell cycle-related pathways, including Myc targets, E2F targets and G2-M checkpoint, and mTORC1 and PI3K/AKT/mTOR signaling, were enriched with downregulated genes. Several pathways related to metabolic reprogramming in cancer cells were also inhibited, i.e. oxidative phosphorylation, glycolysis and hypoxia. Many proinflammatory hallmarks, including TNF-alpha signaling via NF-kB, interferon alpha response, interferon gamma response, IL-2/STAT5 signaling and inflammatory response were enriched with upregulated genes.

### Fulvestrant gene expression signature

(Figure 1D) analysis identified similar enrichment for downregulated genes with estrogen response hallmarks, cell cycle-related pathways, mTORC1 signaling, oxidative phosphorylation and glycolysis evidenced enriched downregulated genes, whereas many proinflammatory hallmarks, EMT, hypoxia and apoptosis were enriched with upregulated genes.

### Endoxifen gene expression signature

(Figure 1C) had common and different traits with those of tamoxifen and fulvestrant. First, endoxifen changed estrogen response genes bi-directionally, with some genes being downregulated, and others being upregulated. The inhibition of cell cycle-related pathways and mTORC1 signaling was detected, but the PI3K/AKT/mTOR pathway was enriched with upregulated genes. Among metabolic reprogramming pathways, glycolysis and oxidative phosphorylation were also changed bi-directionally, but hypoxia and reactive oxygen species (ROS) pathways were strongly upregulated. Finally, apoptosis-related genes were also changed in both directions. These results show that although all three antiestrogens have similar effects, there are substantial differences between them.

### Estrogen-dependent and independent effects of tamoxifen, endoxifen and fulvestrant

The study examined the effect of estrogen signaling on tamoxifen, endoxifen, and fulvestrant. To achieve this, each drug’s signature was intersected with the signature of estrogen and subjected to GSEA analysis.

### Tamoxifen

exhibits both estrogen-dependent and independent mechanisms of action (MOAs). Estrogen-dependent mechanisms predominantly contribute to tamoxifen-mediated downregulation of estrogen response, inhibition of cell cycle-regulating pathways and the promotion of apoptosis. The stimulation of pro-inflammatory pathways by tamoxifen involves both estrogen-dependent and independent MOAs. Tamoxifen’s effects on mTORC1 signaling are mediated through both estrogen-dependent and independent pathways, whereas the regulation of PI3K/AKT/mTOR signaling is mostly estrogen-independent. Among metabolic reprogramming pathways, oxidative phosphorylation is influenced by both estrogen-dependent and independent mechanisms, while hypoxia and glycolysis are mostly estrogen-dependent.

### Endoxifen

is known to have estrogen-dependent and independent effects (Hawse et al. 2013). These analyses further confirmed these data. Dysregulation of estrogen-response pathways was identified as an estrogen-dependent MOA of endoxifen. The modulation of apoptosis, hypoxia and glycolysis by endoxifen was also found to be estrogen-dependent. However, endoxifen-mediated downregulation of cell cycle-regulating pathways was both estrogen-dependent and independent. Both estrogen-dependent and independent mechanisms contributed to endoxifen effects on mTORC1 pathway (Jayaraman et al. 2023). Interestingly, oxidative phosphorylation was enriched with estrogen-independent genes.

### Fulvestrant

demonstrated both estrogen-dependent and independent MOAs. The downregulation of estrogen-response pathways was dependent on estrogen. Both estrogen-dependent and independent mechanisms contribute to fulvestrant’s downregulation of cell cycle-regulating pathways and the promotion of apoptosis. Additionally, both estrogen-dependent and independent genes were enriched with metabolic reprogramming pathways, including oxidative phosphorylation, hypoxia, and glycolysis. However, the stimulation of pro-inflammatory pathways and EMT by fulvestrant are specific to estrogen-independent mechanisms.

### Comparative analysis of endoxifen, tamoxifen, and fulvestrant

Common and drug-specific effects of endoxifen, tamoxifen, and fulvestrant were analyzed by performing comparative analyses by intersecting the gene signature of each agent and identifying shared and unique subsets of genes. GSEA analyses were performed to identify signaling pathways that enrich corresponding gene subsets (Figure 2).

**Figure 2.**
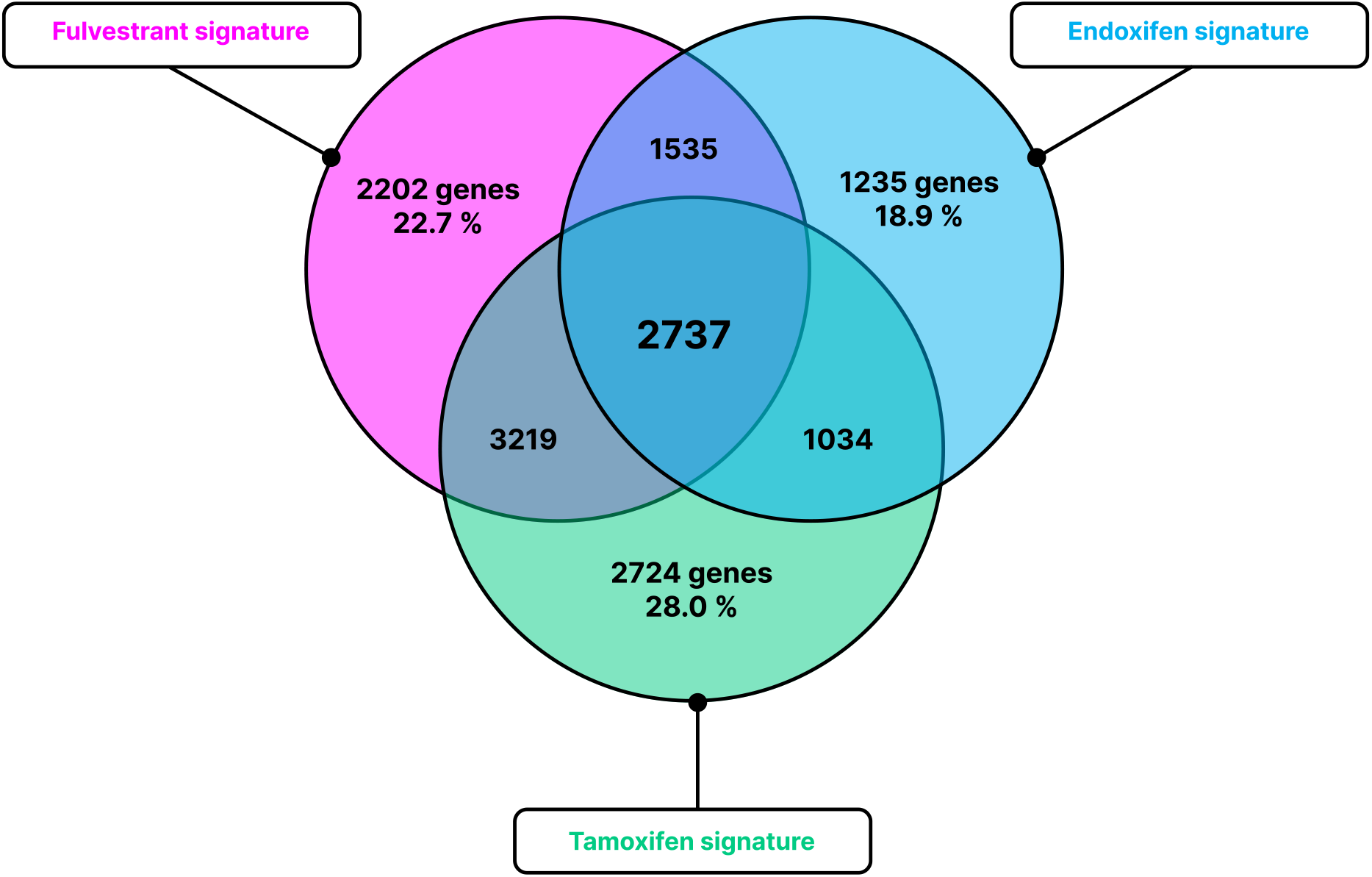
The Venn diagram describes the intersection between genes changed by tamoxifen, endoxifen, and fulvestrant with the indication of numbers of genes in each intersection and the percentages of drug-specific genes among all genes changed by a drug.

2737 genes were identified in the intersection of all three anti-estrogens. There were 1034 genes in the intersection between tamoxifen and endoxifen, 3219 between tamoxifen and fulvestrant, and 1535 between endoxifen and fulvestrant. There were 2724 genes specifically affected by tamoxifen (that accounted for 28.0% of all tamoxifen-dependent genes), 2202 genes specifically affected by fulvestrant (22.7% of all fulvestrant-dependent genes), and 1235 genes specifically affected by endoxifen (18.9% of all endoxifen-dependent genes). Shared and drug-specific gene subsets were analyzed by pathway enrichment analysis. The estrogen signature was included as an additional layer of information for analysis to account for estrogen dependent/independent effects.

The study analyzed the intersection of all three antiestrogens and paired intersections between tamoxifen and endoxifen, tamoxifen and fulvestrant, and endoxifen and fulvestrant. All these gene subsets were characterized by the downregulation of estrogen signaling that was dependent on estrogen because only estrogen-dependent genes in these subsets enriched corresponding hallmarks. The inhibition of cell cycle-related hallmarks and mTORC1 signaling was also a shared effect of all three antiestrogens, which included both estrogen-dependent and independent MOAs. The stimulation of apoptosis was observed in the intersection of the three drugs, and was also observed for the endoxifen/fulvestrant but not for the tamoxifen/fulvestrant intersection. The apoptosis-related genes were mostly estrogen-dependent. Additionally, the activation of pro-inflammatory hallmarks was identified for the overall intersection, and for fulvestrant/tamoxifen but not in fulvestrant/endoxifen intersection. Interestingly, pro-inflammatory effects were estrogen-independent and differed between drugs. Notably, EMT stimulation was typical for fulvestrant/tamoxifen but not for fulvestrant/endoxifen, and was estrogen-independent. Variances in metabolic pathways also appeared in all intersections, and were both estrogen-dependent and independent.

Gene subsets that were specific to each drug were subjected to analysis.

The **Tamoxifen specific gene subset** consisted of 2724 genes, however, early estrogen response was the only hallmark enriched with this subset of genes. Specifically, only estrogen-dependent genes downregulated by tamoxifen were enriched.

The **Endoxifen-specific subset of genes** enriched mTORC1 and PI3K/Akt/mTORC1 pathways. Endoxifen-specific mTORC1 pathway-related genes were both estrogen-dependent and independent, and were either upregulated or decreased after endoxifen treatment. PI3K/Akt/mTORC1 pathway-related genes were mostly estrogen-dependent and were upregulated under the influence of endoxifen.

Although the **fulvestrant specific gene subset** consisted of more than two thousand genes, no enrichment with known hallmarks was identified. This may suggest that the residual fulvestrant-regulated genes are not related to analyzed hallmarks.

### Analysis of potential molecular effects of tamoxifen, endoxifen and fulvestrant in breast cancer

The analysis of gene expression changes in the MCF7 cell line identified the alterations induced by tamoxifen, endoxifen, and fulvestrant in breast cancer cells. To determine the relative effect of these alterations on breast cancer, a molecular signature for breast cancer was created comprising the most significantly altered genes, validated across multiple omics datasets integrated in PandaOmics (Figure 3A). Each drug signature was compared with the disease signature, focusing on gene sets exhibiting opposite expression patterns. Specifically, genes were identified that are upregulated in breast cancer but downregulated after treatment. Conversely, genes were as well identified that are downregulated in cancer compared to healthy tissue but upregulated following treatment. GSEA analyses were performed on the identified gene subsets for each drug.

**Figure 3.**
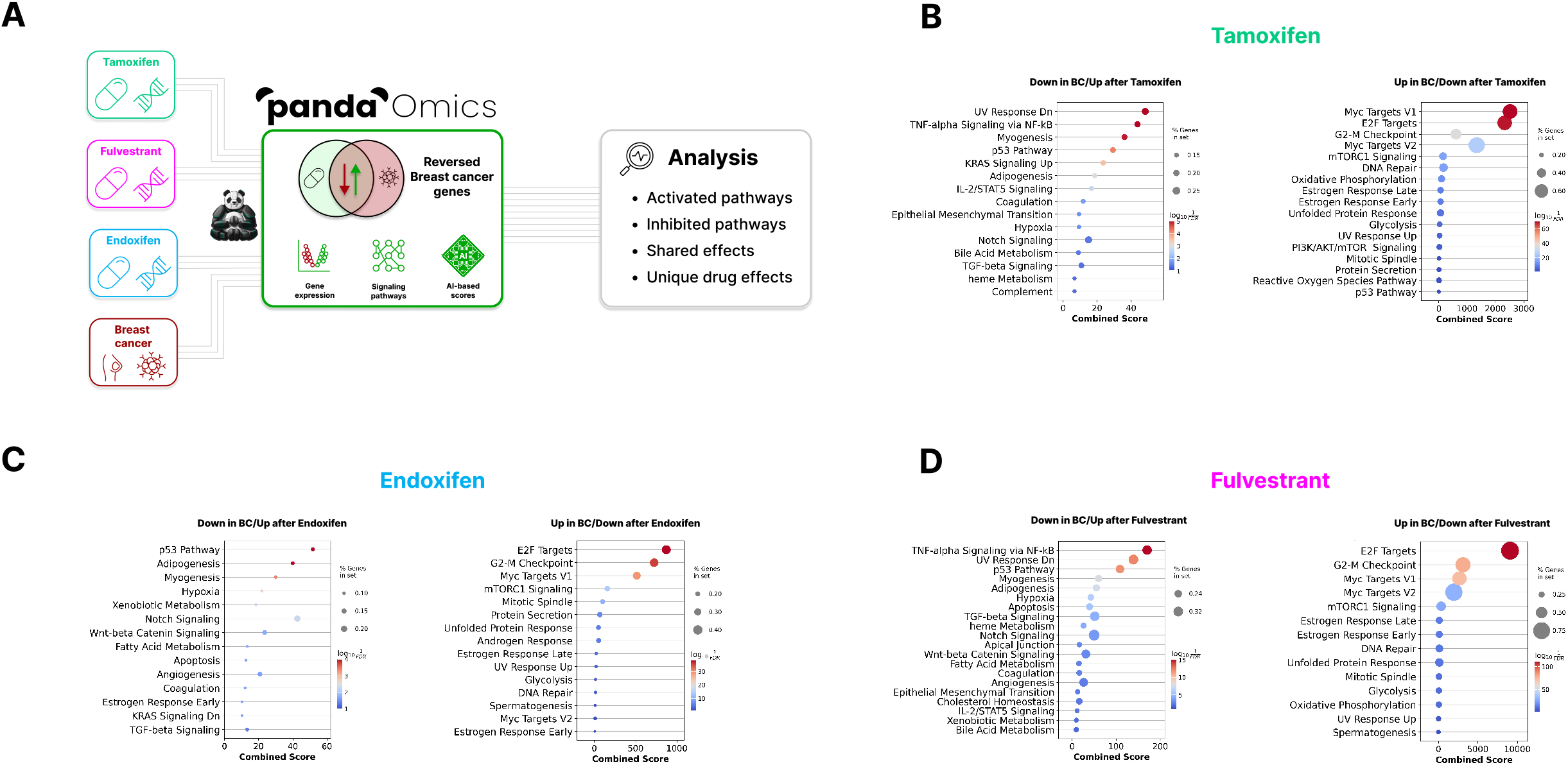
A) Breast carcinoma analysis pipeline. Breast carcinoma molecular signature was developed based on multiple omics datasets integrated in PandaOmics. Each drug-specific gene signature was compared with the disease signature. GSEA analysis was conducted on gene sets exhibiting opposite expression patterns in tamoxifen (B), endoxifen (C) or fulvestrant (D) treated cells and in breast carcinoma. Inhibited and activated pathways that are shared between all three drugs or are specific for one or two drugs were identified.

The results of these analyses are presented in Figure 3. As expected, a subset of estrogen signaling-related genes upregulated in breast carcinoma were inhibited by all three anti-estrogens, underscoring their efficacy in suppressing estrogen signaling in breast cancer cells. Notably, these analyses revealed the presence of early estrogen response genes that are inhibited in breast carcinoma but which can be restored by endoxifen, indicating a unique bi-directional regulation of the estrogen response by endoxifen, which has not been previously reported.

All three drugs significantly downregulated cell cycle hallmark genes that are commonly hyperactivated in breast carcinoma, reflecting their antiproliferative effects observed in vitro and in vivo. Interestingly, apoptosis-related genes downregulated in breast carcinoma were reversed by endoxifen and fulvestrant but not by tamoxifen. Both tamoxifen and fulvestrant stimulated a subset of genes related to the EMT pathway that were decreased in breast carcinoma. Similarly, the stimulation of pro-inflammatory IL-2/STAT5 signaling and TNF-alpha/NFkB pathways in breast cancer was common to tamoxifen and fulvestrant but not to endoxifen.

Among metabolic reprogramming pathways, hypoxia-enriched genes downregulated in breast cancer were upregulated by all three antiestrogens, while glycolysis-enriched genes upregulated in breast cancer were downregulated by all three anti-estrogens. Additionally, genes upregulated in breast cancer and downregulated by tamoxifen or fulvestrant were enriched with oxidative phosphorylation, whereas only tamoxifen-regulated genes were enriched with ROS pathway.

## Discussion

Breast cancer remains one of the most prevalent malignancies affecting women worldwide, with estrogen receptor-positive (ER+) breast cancer constituting a significant proportion of these cases (Yadav et al. 2024). Historically, endocrine therapy, including tamoxifen, fulvestrant and aromatase inhibitors, has been the mainstay of treatment for ER+ breast cancer. Tamoxifen belongs to a class of SERMs that interacts with estrogen receptors in a tissue-specific manner, acting as estrogen agonists or antagonists depending on the target tissue (Maximov, Lee, and Jordan 2013). Unlike tamoxifen, fulvestrant is a SERD, primarily targeting the estrogen receptor alpha, leading to its degradation and subsequent inhibition of estrogen-dependent transcription and cell proliferation (Patel et al. 2023). However, in addition to varying efficacy across different breast cancer subtypes, the development of resistance to endocrine therapy poses a substantial challenge (Jeselsohn et al. 2017). Moreover, vasomotor symptoms such as hot flashes are often reported in tamoxifen-treated patients, leading to reduced quality of life and hampering patients’ daily functioning (Chan et al. 2021). Importantly, HRT use at clinical trial entry or during its course was not effective in alleviating hot flashes for women treated with tamoxifen (Sestak et al. 2006). All of the above indicates the need to explore additional therapeutic strategies.

Endoxifen, identified as the predominant active metabolite of tamoxifen, functions as a potent anti-estrogenic compound, inhibiting the growth of various breast tumor cell lines and being rapidly absorbed when administered orally (Jayaraman et al. 2021; Ahmad et al. 2010). Emerging studies have focused on endoxifen and demonstrated endoxifen’s efficacy and safety, suggesting that it could be a valuable alternative in the treatment of ER+ breast cancer due to its potent anti-estrogenic effects and dual MOA with PKCβ inhibition (Goetz et al. 2017; Jayaraman et al. 2020).

This study provides useful insights into the MOA of endoxifen compared to tamoxifen and fulvestrant and an analysis of their potential effects in breast cancer. Artificial intelligence tools generally have made positive contributions in the prediction of molecular expression (Yousif et al. 2022). The AI-driven PandaOmics platform was utilized in conjunction with bioinformatics approaches to construct gene signatures of the three drugs and identify key pathways determining their anticancer effects. These findings highlight the differentiation of endoxifen compared to tamoxifen and fulvestrant, due to endoxifen’s distinct MOA.

Firstly, utilizing a computational approach, individual molecular signatures were developed for tamoxifen, endoxifen, and fulvestrant in the MCF7 cell line, a well-established and widely used human ER+ breast cancer model. These signatures were subjected to independent analysis and also to a comparative analysis conducted by intersecting them and examining individual gene sets through a Venn diagram analysis.

Secondly, potential tamoxifen, endoxifen, and fulvestrant-regulated pathways and downstream mediators were identified that are dependent or independent on estrogen signaling. This was achieved by intersecting the molecular signature of each agent with that of estrogen. All three anti-estrogens were observed to exert both estrogen-dependent and independent effects. Substantial evidence from clinical trials has demonstrated efficacy of tamoxifen and hormone therapy in ER negative (ER-) cancers (Kirkby et al. 2023; Scarpetti et al. 2023; Yang et al. 2012). These analyses and results therefore offer evidence of pathways regulated by endoxifen, tamoxifen or fulvestrant that may underlie the efficacy of these drugs in ER negative cancers.

Thirdly, key pathways dysregulated in breast cancer patients were identified that can be oppositely modulated by anti-estrogens. To achieve this, a molecular signature for breast cancer was developed and intersected with the drug signatures, focusing on gene sets displaying opposite expression patterns in the disease compared to post-treatment with anti-estrogens.

The data indicated that all three anti-estrogens significantly downregulated estrogen signaling. Importantly, the variances in the estrogen response hallmarks were strictly estrogen-dependent, validating these data and analyses. Moreover, the analysis of breast carcinoma omics data identified a subset of estrogen response genes that are upregulated in breast carcinoma but inhibited in all three anti-estrogen signatures, highlighting the effective role of anti-estrogens in shutting down estrogen signaling in breast cancer (Miziak et al. 2023). It should be noted that SERMs exhibit bi-directional effects on estrogen receptor signaling by acting as either agonists or antagonists depending on the tissue context and the specific ER isoforms involved. Importantly, current analyses also revealed early estrogen response genes that are inhibited in breast carcinoma but which can be upregulated by endoxifen but not by tamoxifen or fulvestrant. Endoxifen is known to have concentration-dependent effects on estrogen signaling (Hawse et al. 2013; Jayaraman et al. 2023), but these data are the first report on the bi-directional character of these effects. We hypothesize that endoxifen’s bi-directional modulation of early estrogen response genes may have beneficial clinical outcomes in breast cancer. Specifically, certain early estrogen response genes, which are upregulated by endoxifen and downregulated in breast carcinoma, are known to possess antitumorigenic properties in breast cancer. For example, the *IGFBP4* gene encodes Insulin-like Growth Factor Binding Protein 4, which is upregulated by estrogen receptor alpha and acts as an anti-cancer protein in ER+ breast cancer, correlating with improved survival (Mita et al. 2007; Flynn and Houston 2022). The *KRT15* gene, stimulated by endoxifen, is lower in breast carcinoma than normal tissues and is associated with better survival and immune cell infiltration, varying across breast cancer subtypes (Zhong et al. 2021; Zhang et al. 2023). *OLFML3* low expression is detected in the neoadjuvant chemotherapy resistant group of breast cancer patients and is associated with low overall survival pointing to its antitumorigenic role (Barrón-Gallardo et al. 2022). *PEX11A* expression is linked to changes in the immune-infiltrating microenvironment after paclitaxel treatment and correlates with better survival (Shen et al. 2020). THSD4, involved in extracellular matrix assembly, is downregulated in breast cancer, and high expression predicts better prognosis (Bao et al. 2022). These findings highlight endoxifen’s potential in activating beneficial estrogen response genes in breast cancer. Similarly, tamoxifen’s influence extends beyond its anti-estrogenic activity in breast tissue. For example, tamoxifen’s pro-estrogenic effects demonstrated beneficial effects on muscle pathology, particularly in the context of Duchenne Muscular Dystrophy (DMD) (Dorchies et al. 2013), and on bone, which has made it attractive for the treatment of osteoporosis (Jordan 2008; Singh, Martin-Hirsch, and Martin 2008).

All three studied anti-estrogens downregulated cell cycle-related genes, which includes both estrogen-dependent and independent mechanisms of action. This is in agreement with previous reports on endoxifen from *in vitro/in vivo* evidence, supported by transcriptomics analysis which underscores the potent antitumor and antiproliferative activity of endoxifen (Jayaraman et al. 2020; Helland et al. 2015).

The activation of pro-inflammatory hallmarks was identified for all three anti-estrogens. However, it was more pronounced for tamoxifen and fulvestrant, especially in the context of breast carcinoma signature. The activation of the pro-inflammatory response suggests potential strategies for combination therapies with immune checkpoint inhibitors (ICIs) in ER+ breast cancer. This is particularly relevant given the limited efficacy of ICIs in hormone receptor (HR)-positive breast cancer, which is attributed to the low tumor mutational burden, low percentage of stromal tumor-infiltrating lymphocytes (sTIL), and low PD-L1 expression (Li, van der Merwe, and Sivakumar 2022). Conflicting evidence has been reported for fulvestrant with ICIs, suggesting that the immune-modulatory effects of fulvestrant may vary depending on the treatment duration, dose, and presence of additional factors such as IFN-gamma (Hühn et al. 2022; Hermida-Prado et al. 2023; Huang et al. 2021). A phase II study assessing efficacy and safety of combination of nivolumab (ICI inhibitor) with abemaciclib (a CDK4/6 inhibitor) and endocrine therapy (either fulvestrant or letrozole) in patients with HR-positive, HER2-negative metastatic breast cancer was terminated early due to safety concerns. While the combination therapy was active, it was associated with significant immune-related hepatotoxicity (Masuda et al. 2023). These findings, taken together with this study, highlight the potential of endocrine therapy to augment immune responses but underscore the need for strategies to manage proinflammatory cytokine production and associated immune-related adverse events. It is worth noting that this analysis utilized transcriptomic data from the MCF7 cell line, which does not fully represent the complex immune environment. Using transcriptomic data from more complex experimental models, including various immune cell types, would provide a more accurate assessment of the effects of the three drugs on pro-inflammatory signaling pathways.

The PI3K/AKT/mTOR pathway is a critical driver implicated in breast cancer, often associated with resistance to endocrine therapies (Miricescu et al. 2020). mTORC1 is a specific complex within this pathway that plays a pivotal role in regulating cell growth and metabolic processes. The mTORC1 and PI3K/AKT/mTOR hallmarks overlap to some extent, however they represent different subsets of genes up-regulated through activation of the mTORC1 complex or by activation of the PI3K/AKT/mTOR pathway (Liberzon et al. 2015). It was notable that while downregulated genes in all three anti-estrogen signatures were enriched with the mTORC1 pathway, endoxifen-upregulated genes were specifically enriched with the PI3K/AKT/mTOR pathway. Furthermore, analysis of a subset of endoxifen-specific genes, unaffected by either tamoxifen or fulvestrant, revealed enrichment of both mTORC1 and PI3K/AKT/mTORC1 pathways. No similar enrichment was found for tamoxifen- or fulvestrant-specific subsets. Recent studies have demonstrated that endoxifen downregulates AKT phosphorylation through protein kinase C beta 1 (PKCβ1) inhibition, and suggested that proteins MTOR, RPS6K, and AKT1 involved in the AKT signaling pathway could potentially be regulated by high-dose endoxifen (Jayaraman et al. 2023). Our findings corroborate the modulation of the PI3K/AKT/mTOR pathway by endoxifen, and suggest that this effect is more specific to endoxifen than tamoxifen or fulvestrant.

Stimulation of the EMT was observed in fulvestrant signature and its intersection with tamoxifen, and was independent of estrogen signaling. A specific subset of genes associated with the EMT pathway was identified that was downregulated in breast carcinoma but stimulated by both tamoxifen and fulvestrant, while not being affected by endoxifen. This implies that tamoxifen and fulvestrant exert distinct effects on EMT-related genes, potentially contributing to the unique mechanisms of action observed in breast cancer.

The stimulation of apoptosis was observed in the gene signatures of fulvestrant and endoxifen, but not tamoxifen. The intersection of all three drug signatures exhibited apoptosis stimulation, which was also typical for the endoxifen/fulvestrant intersection but not for the tamoxifen/fulvestrant intersection. Most apoptosis-related genes were estrogen-dependent. Interestingly, genes related to apoptosis that were downregulated in breast carcinoma were reversed by endoxifen and fulvestrant, but not by tamoxifen. Consistently, recent studies demonstrate that endoxifen treatment results in induction of apoptosis through the suppression of AKT signaling in ERα+ breast cancer cells (Jayaraman et al. 2023). These data may indicate a more pronounced role for endoxifen in promoting apoptosis compared to tamoxifen.

Metabolic reprogramming is a known driver of cancer (Sebastian 2014) and dysregulation of pathways involved in metabolism was determined in the analyses of this study. Both oxidative phosphorylation and glycolysis pathways were enriched with genes downregulated by tamoxifen and fulvestrant. However, endoxifen affected these pathways bi-directionally, with enrichment observed in both upregulated and downregulated genes. Furthermore, ROS-related genes were exclusively upregulated by endoxifen. Analysis of the breast cancer signature revealed subsets of metabolic genes that were oppositely regulated by antiestrogens. Genes downregulated in breast cancer but upregulated by all three antiestrogens were enriched for the hypoxia hallmark, whereas genes upregulated in breast carcinoma but inhibited by all three antiestrogens were enriched for glycolysis. Additionally, genes upregulated in breast cancer and downregulated by tamoxifen or fulvestrant were enriched for oxidative phosphorylation, while only tamoxifen-downregulated genes were enriched for the ROS pathway. This differential regulation of metabolic pathways by antiestrogens underscores their unique mechanisms of action in breast cancer.

In conclusion, this study utilizing AI-driven and bioinformatics approaches aimed to uncover the MOA of endoxifen compared to tamoxifen and fulvestrant. The findings made in this study highlight differentiation in the molecular signature for endoxifen compared to its parent compound tamoxifen, in addition to shared effects with tamoxifen or fulvestrant. However, further research is needed to advance the understanding of endoxifen’s MOA. Experimental validation of identified pathways in cell lines and animal models is crucial to confirm the mechanistic insights and therapeutic potential of endoxifen.

One limitation of the current study is the lack of comprehensive proteomics data. One proteomic dataset was available for tamoxifen and two for breast cancer, but not for the other signatures. Consequently, these findings are primarily based on mRNA changes rather than protein levels. Translating these data to the protein level poses challenges due to discrepancies often observed between proteomic and transcriptomic data (Prabahar et al. 2024). Moreover, mRNA and protein levels do not always correlate, partly due to internal adaptive mechanisms such as translational offsetting, which can regulate protein levels independently of mRNA changes. This mechanism is particularly described for estrogen-regulated transcripts (Lorent et al. 2019) and so may have a strong effect upon the analyses of endoxifen, tamoxifen and fulvestrant. Nonetheless, this study provides support for the use of transcriptomic data to analyze drug mechanisms and the belief that integrating multi-omics data, including proteomics and metabolomics, could offer a more comprehensive understanding of their molecular effects. A more comprehensive understanding of the molecular effects of drug mechanisms would provide benefits with respect to determining the optimal application of therapeutics for existing indications. In addition, if this use of transcriptomic data is successful, it could be expanded to find potential novel, repurposed uses of existing therapeutics in novel indications as validated subjects for study.

## Authors contribution

AS conducted and coordinated *in silico* studies, AV, MK designed and conducted *in silico* analysis and wrote the manuscript, KMA contributed to the writing of the manuscript. AZ and SQ supervised the project and provided suggestions to the design and improvement of the manuscript. SSH and HS provided helpful suggestions and contributed to the writing of the manuscript, HLR contributed to writing the manuscript and is the corresponding author.

## Data availability

All data supporting the conclusions of the paper are available in the article and corresponding figures. Datasets used in the paper are described in the materials and methods section.

